# Colourful predictive templates in early visual cortex

**DOI:** 10.64898/2026.06.11.731729

**Authors:** Mandy Viktoria Bartsch, Eelke Spaak, Floris Pieter de Lange

**Affiliations:** Donders Institute for Brain, Cognition, and Behaviour, Radboud University, 6500 GL Nijmegen, The Netherlands

## Abstract

Predictive processing theories suggest that our brain constantly predicts incoming sensory information, though the level of granularity and concrete sensory content of these predictions is debated. In this study, we investigate whether predictions in visual cortical areas carry information about colour before the onset of the anticipated stimulus. Thirty-seven human participants of both sexes viewed coloured disc stimuli presented in a predictable sequence, enabling the brain to form expectations about the upcoming colour, while the brain response was tracked with high spatiotemporal resolution (combined electro- and magnetoencephalography). A decoding model trained on a separate colour localizer dataset could successfully reconstruct the predicted colour during the pre-stimulus period, where only a grey placeholder was present. Thereby, our findings demonstrate that early visual regions of the brain form anticipatory representations of colour information, in the absence of external colour input, yet in a representational format similar to veridical sensory input.

**Significance Statement:** The human brain continuously anticipates upcoming events, but the exact features and level of detail of these predictions is a topic of debate. Here we show that when a visual stream has a predictable sequence of colours, the brain pre-activates colour information in early visual cortex before the stimulus appears. Using high-temporal-resolution brain recordings and multivariate decoding analyses, we demonstrate that expected colours can be reconstructed from neural activity several hundreds of milliseconds before the anticipated stimulus is shown. These findings provide direct evidence that perceptual predictions contain concrete sensory content in terms of colour information. By clarifying the features that make up predictive representations, our work advances understanding of how expectations shape perception and guide attention in everyday vision.

## Introduction

Whether catching a ball or navigating city traffic, the brain must constantly predict what will happen next. Because visual processing and motor responses take time, the brain relies on statistical regularities and prior experience to anticipate events—such as a ball’s trajectory or changing colours of traffic lights. While predictive processing theories suggest that the brain continuously generates predictions about incoming sensory input (Friston, 2010; Rao and Ballard, 1999), the specific form of these predictions is less clear: Do we construct detailed sensory representations of the expected stimuli, or do predictions remain more abstract?

We address this question for a fundamental visual feature: colour. Colour is central to perception, as it guides scene segmentation (Tanaka et al., 2001), facilitates face (Yip and Sinha, 2002) and object recognition (Bramão et al., 2011), supports scene memory (Spence et al., 2006; Wichmann et al., 2002), and is a powerful guiding cue in visual search (Wolfe and Horowitz, 2004). Given its pervasive role in perception, attention, and memory, predictive mechanisms may also incorporate colour information.

An elegant way to uncover the feature content of internally generated expectations is to examine brain activity in the absence of visual input, thereby isolating prediction signals from sensory-driven responses. This can be accomplished by decoding neural responses to omitted or occluded stimuli (Ekman et al., 2017; Erlikhman and Caplovitz, 2017; Kok et al., 2013; Papale et al., 2023; Smith and Muckli, 2010), or from pre-stimulus activity in cueing designs (Gong et al., 2022; Kok et al., 2017; van Moorselaar and Slagter, 2019).

Building on these approaches, recent fMRI studies have demonstrated prediction-related activity in early visual areas, suggesting that expectations about occluded greyscale scenes are represented in a coarse structural visual format (Morgan et al., 2019). Researchers have also decoded expected motion trajectories (Ekman et al., 2017) and anticipated stimulus orientations from omission trials (Kok et al., 2013), with such predictions likely originating in deep layers of V1 (Aitken et al., 2020). High temporal resolution MEG further revealed stimulus-specific orientation patterns emerging ∼40 ms before stimulus onset, reflecting a pre-stimulus expectation template for orientation (Kok et al., 2017). Together, these studies indicate that early visual cortex carries visuo- spatial/retinotopic information about anticipated input. However, a key gap in our understanding of sensory predictions is whether they also encode non-spatial features such as colour.

As a necessary foundation for this question, previous studies have established the ability to decode colour information from non-invasive recordings. fMRI studies demonstrated reliable colour decoding in early visual areas (V1–V3) and ventral stream regions V4 and VO1/2 (Brewer et al., 2005; Brouwer and Heeger, 2013, 2009), with colour representations becoming more perceptually aligned in V4 and VO1. More recent EEG (Chauhan et al., 2023; Sutterer et al., 2021) and MEG studies (Hermann et al., 2022; Rosenthal et al., 2021) could decode colour information with high temporal resolution, with early visual areas (V1-V4) likely driving the EEG and MEG colour signals (Sandhaeger et al., 2019). In addition, colour information stemming from object knowledge or memory representations has been successfully decoded from responses to greyscale images (Bannert and Bartels, 2013; Retter et al., 2023; Teichmann et al., 2019), and after stimulus offset (Bae and Chen, 2024; Bocincova and Johnson, 2019; Hajonides et al., 2021; Kandemir et al., 2024), opening up the question whether the mere anticipation of a colour activates colour-tuned sensory representations.

In the current study, we used concurrent EEG/MEG and multivariate pattern analysis (MVPA) to examine whether colour expectations in visual cortex form before stimulus onset when only a grey placeholder was presented. To elicit colour expectations, coloured discs were shown in a predictable sequence while participants performed an orthogonal hue-change detection task. A decoding model trained on perceptual colour responses could successfully reconstruct both post-stimulus perceived colour and pre-stimulus expected colour. These results show that predictions lead to the formation of colourful sensory templates in early visual cortex prior to stimulus onset.

## Materials and Methods

### Participants

Fourty-five healthy adult volunteers participated in the study, out of which eight had to be excluded due to technical difficulties, poor data quality (excessive blinking, tiredness, noisy sensors), or incomplete datasets, leaving thirty-seven participants (mean age 23.6 years, 10 male, 3 left-handed). All participants had normal or corrected-to-normal visual acuity and normal colour vision, as assessed using Ishihara plates (Test Chart Books for Color Deficiency, 24 plates). Participants were recruited via word-of-mouth and from the student population through the electronic SONA system. Written informed consent was obtained from all participants prior to the experiment, and they received monetary compensation for their participation (total duration, including preparation and measurement, 3–4 h). All experimental procedures were approved by the local ethics committee (CMO region Arnhem-Nijmegen, The Netherlands) and conducted in accordance with the general ethics approval (CMO2014/288, “Imaging Human Cognition”). Trial numbers and sample size were based on previous studies on visual expectation (Kok et al., 2017, 2014), colour tuning (Bartsch et al., 2017; Schulz et al., 2024), and colour decoding experiments (Hajonides et al., 2021; Hermann et al., 2022; Sutterer et al., 2021), sample size was according to G*Power calculations (RRID:SCR_013726) (Faul et al., 2007) suited to detect an effect with 80% power, assuming effects of medium effect size or larger (Cohen’s d>0.5, significance level 0.05, within-subject design, paired t-test).

### Experimental Design

#### General Paradigm

We examined whether brain activity patterns in early visual cortex contained information about (expected) stimulus colour, both after and before stimulus onset. The experimental design is depicted in Figure 1. To measure sensory responses to colour, participants were presented with coloured annuli (ring-shaped stimuli) while performing a task at fixation (colour localizer blocks). In separate experimental blocks, colour expectations were induced by presenting the same stimuli in a structured sequence with 80% validity (colour expectation task). Because the formation of colour expectations likely depends on attention to the coloured stimuli (Richter and de Lange, 2019), participants were instructed to detect occasional hue changes of the annuli. Notably, this task was designed to be orthogonal to the predictive manipulation (i.e., colour order), ensuring that expectation-related effects were not confounded by task relevance (e.g., preparatory motor responses or colour naming; see also Discussion).

**Figure 1.**
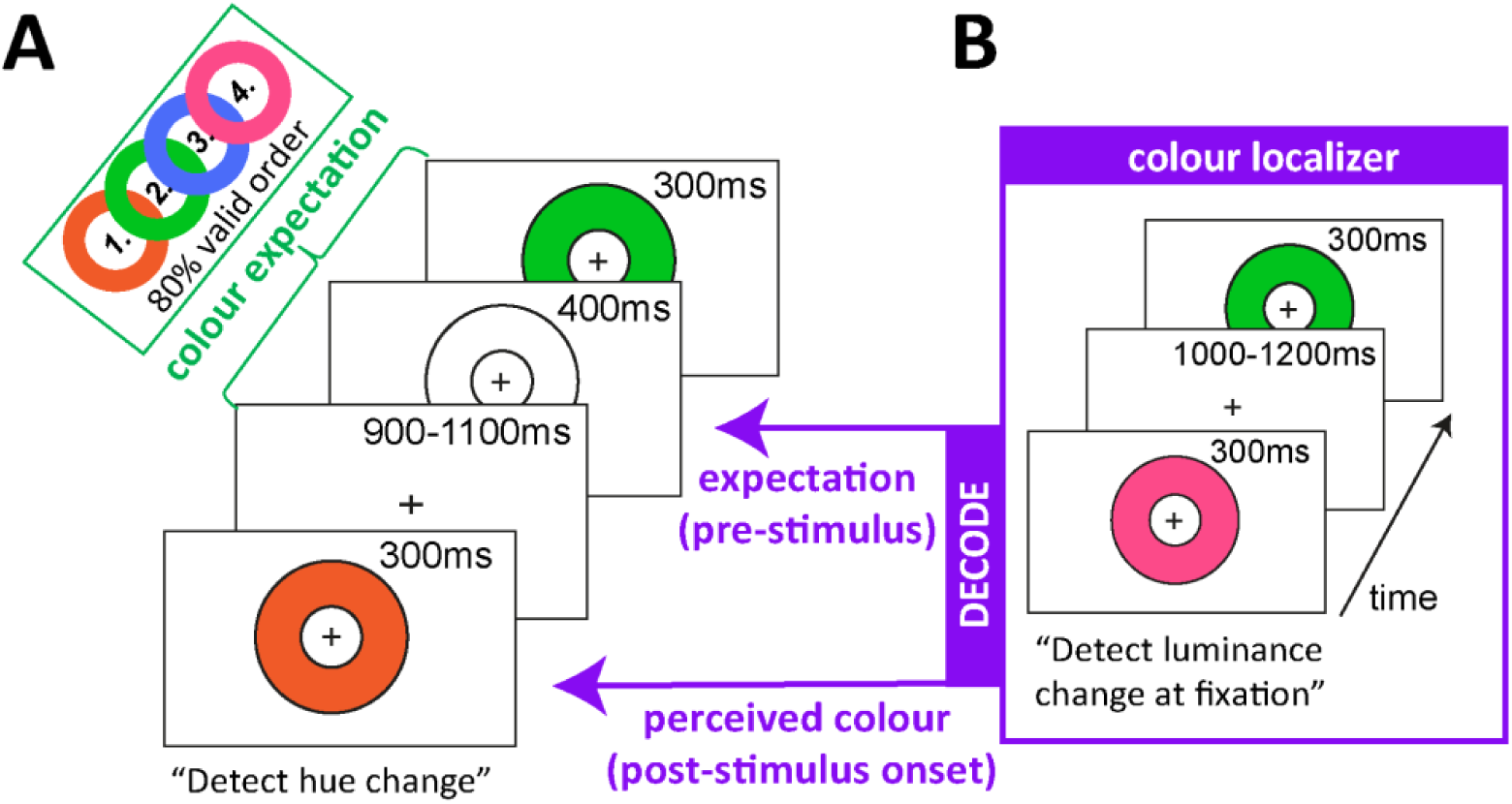
Experimental Design. **A)** Colour Expectation task. Participants were shown coloured annuli (ring-shaped stimuli) in an 80% valid sequence while detecting a transient hue change (e.g., orange becoming yellowish) of the currently presented annulus. The coloured annuli were preceded by a grey outline, which served as a temporal cue to facilitate the formation of expectations. **B)** Colour localizer runs. Participants attended to luminance changes of the fixation cross while coloured annuli were presented in random order. That way, stimuli were task-irrelevant and elicited purely sensory colour responses, without involving processes of discrimination, decision-making, or colour-naming. A decoding model based on these sensory colour responses was then used to isolate cortical colour representations from the expectation task, both post-stimulus onset (reflecting perceived colour, later referred to as “stimulus decoding”), and pre-stimulus onset during the colour expectation phase, when only the grey placeholder was on the screen (later referred to as “template decoding”).

**Figure 2.**
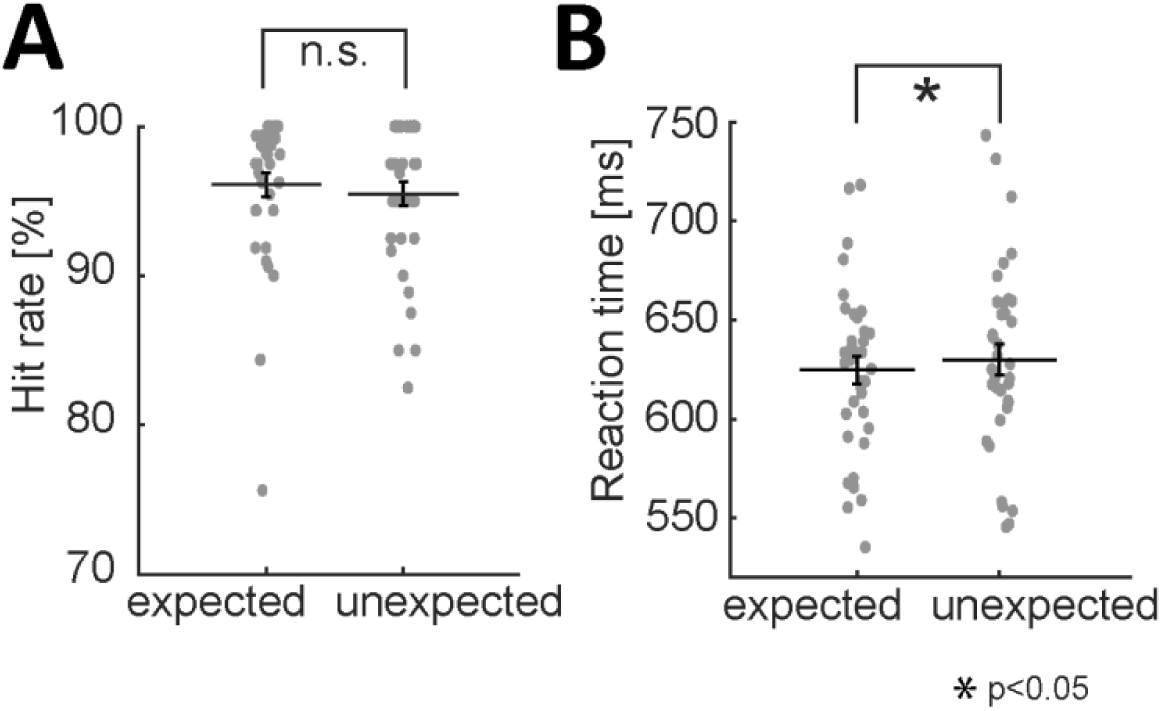
Behavioural performance on hue change trials in colour expectation blocks. For both panels, black horizontal lines represent the mean, error bars show the standard error of the mean (SEM). Grey dots depict performance of individual participants. **A)** Hit rate was close to ceiling for most of the participants regardless of colour sequence expectation. **B)** Reaction times were slightly faster for expected compared to unexpected trials.

#### Stimuli

Stimuli are illustrated in Figure 1. A coloured annulus was presented concentrically around a central fixation cross, with an inner radius of 1.1° and an outer radius of 3.6° of visual angle. Colours were defined in DKL space (Derrington et al., 1984) and presented on an isoluminant grey background (82 cd/m²) at a viewing distance of 80cm. Based on previous work on colour decoding using MEG (Rosenthal et al., 2021), the four hues were selected along the intermediate axes of DKL space: 45° (magenta), 135° (blue), 225° (green), and 315° (orange).

In the localizer runs, stimulus colour varied pseudo-randomly across trials, with the constraint that the same colour was not repeated on consecutive trials. On 10% of the trials, the fixation cross briefly increased in luminance (100 ms), serving as a target event. These events could occur either during stimulus presentation or in the interstimulus interval (ISI), which ranged from 1000 to 1200 ms (rectangular distribution).

In the expectation task, ISIs ranged from 900 to 1100 ms (rectangular distribution). Here, colours were presented in an 80% valid sequence, such that each participant was assigned one of four predefined cyclic colour orders (e.g., 1–2–3–4 and permutations thereof). For each participant, the assigned sequence was reversed halfway through the expectation blocks. The presentation of the coloured annulus was preceded by a dark grey outline of the stimulus (9.4 cd/m²) presented 400 ms before the colour was filled in. This placeholder served as a temporal cue to facilitate the formation of expectations and to promote the allocation of internal attention to the upcoming stimulus (Bartsch et al., 2018; Jefferies and Di Lollo, 2015; Wildegger et al., 2017; Woodman et al., 2009). On 10% of trials, the hue of the stimulus changed 100 ms after onset, transitioning gradually over 200 ms by 35° in DKL space (i.e., orange and green shifted toward yellow, whereas magenta and blue shifted toward purple).

#### Procedure

Localizer blocks and the expectation task were performed within the same MEG session. Participants first completed three colour localizer blocks (4.5 min each; 160 trials per block), before colours or their order gained relevance. This design allowed us to isolate sensory responses to colour independent of later expectation or colour discrimination processes. Subsequently, participants performed ten blocks of the expectation task (6.5 min each; 200 trials per block). Before the main expectation task, participants completed a short training block (∼2 min; 60 trials) to familiarize themselves with the hue-change detection task and to implicitly train the colour sequence, which was 100% valid during training. After five experimental blocks, the colour sequence was reversed, and participants completed an additional training block (∼2 min) with the new sequence. Participants were informed about the colour order at the beginning of each block (“colours will be mainly presented in the following order: …”), though most of them reported paying no or only some transient initial attention to it, with some of them concluding after encountering invalid colour order trials that there is in fact no such sequence. Trials containing fixation cross luminance changes (localizer) or hue changes (expectation task) were excluded from analysis. This resulted in 108 trials per colour in the localizer and 450 trials per colour in the expectation task, of which 360 corresponded to validly predicted stimuli (expected colour presented) and 90 to invalidly predicted stimuli (unexpected colour presented). In all experimental blocks, participants were instructed to maintain fixation on the central cross and minimize eye blinks. To facilitate this, brief blinking pauses (7 s) were provided approximately every 30–40 s.

### Data acquisition

EEG and MEG were simultaneously recorded. Participants were equipped with an electrode cap with MEG-compatible mounted sintered Ag/AgCl electrodes (Easycap, Herrsching, Germany), and seated in a dimmed, magnetically shielded recording booth (μ-metal; Vacuumschmelze, Hanau, Germany) below the MEG dewar in front of a partly transparent screen at a viewing distance of 80cm. Stimuli were back-projected onto the screen using an DLP-LCD projector (ProPixx, VPixx Technologies Inc., Saint-Bruno, QC, Canada; resolution: 1920 × 1080; refresh rate: 120 Hz, RRID:SCR_013299) placed outside the booth. Stimuli were delivered using MATLAB (The Mathworks, Inc., Natick, MA, USA; RRID:SCR_001622) Psychtoolbox software and custom-written scripts. Participants gave responses with their index finger of the right hand using an MEG compatible handheld button box. Eye position and pupil size were recorded with an SR Research Eyelink 1000 eye tracker, the signal was fed as additional channels in the MEG recording. EEG and MEG were low-pass filtered online at 300 Hz and digitized at a sampling rate of 1200 Hz.

#### EEG recording

The electroencephalogram (EEG) data were continuously recorded using a 32-electrode cap with mounted sintered Ag/AgCl electrodes (Easycap, Herrsching, Germany). Electrode positions were chosen according to the international extended 10-20-system (American Electroencephalographic Society, 1994): Fp1, F7, F3, FC1, C3, CP1, T7, P7, Fp2, F8, F4, FC2, C4, CP2, T8, P8, Fz, Cz, Pz, Oz, Iz, P3, PO7, PO9, PO3, O1, P4, PO8, PO10, PO4, O2. Fpz served as ground. Contact between electrodes and head surface was established using an initial scrubbing with alcohol followed by applying the abrasive electrolyte gel Abralyt light (Easycap, Herrsching, Germany). Impedances were kept below 5kΩ for all electrodes. An electrode at the right mastoid served as online reference during recording. Data were offline re-referenced to the weighted mean of the electric activity of the electrodes at the left and right mastoid according to (Luck, 2005, pp. 107– 108).

#### MEG recording

The magnetoencephalogram (MEG) was recorded using a whole-head 275-channel axial gradiometer CTF MEG system (CTF MEG systems). Head position of the participants was monitored online with three localizer coils placed at the left and right ear as well as the nasion. Deviations of more than 5mm were corrected in between experimental blocks using a visual tool where participants can see their own head position and self-reposition (Stolk et al., 2013). To co-register anatomical and MEG data for source localization, the head shape and the three localizer coils were digitized using a Polhemus 3D tracking device (Polhemus, Colchester, VT, USA).

#### MRI scans

In a separate session, T1-weighted anatomical MRI scans were acquired on a 3T MRI system (Siemens) using an MP-RAGE sequence (GRAPPA acceleration factor of 2, TR = 2.3 s, TE = 3.03ms, voxel size 1 mm isotropic, 192 sagittal slices, 8 ° flip angle). For co-registration with MEG, Vitamin E pills were placed at the left and right ear as well as the nasion.

### Data analysis

#### Behavioural data

Temporal onsets of stimuli and response times were derived from logfiles generated with MATLAB (MathWorks Inc., Natick, MA, USA; RRID:SCR_001622). Hit rates and false alarm rates were computed separately for the luminance detection task (localizer blocks) and the hue change detection task (expectation blocks). Reaction time analyses were restricted to correctly detected targets (hits). For statistical validation, paired Student’s t-tests were used as implemented in MATLAB with an alpha of 0.05 serving as significance level. For an exploratory post hoc analysis (Supplementary Material, Figure S1), participants were categorized based on whether they exhibited numerically faster reaction times for hue changes in expected versus unexpected colours. Participants showing such a pattern were classified as demonstrating a reaction time benefit (“RT benefit expected”, *n* = 25), whereas the remaining participants were classified as showing no benefit (“no RT benefit”, *n* = 12).

#### EEG / MEG – epoching, artefact rejection, event-related fields

MEG and EEG data were analysed offline using the FieldTrip toolbox in MATLAB (Oostenveld et al., 2011) (RRID:SCR_004849). The continuous data were epoched offline relative to the onset of the coloured stimulus from -0.25 to 0.65 s (localizer) and -0.65 to 0.65 s (expectation task). Data were demeaned and line noise was attenuated using a discrete Fourier transform filter at 50 Hz and its harmonics (up to 250Hz). Trials and channels were first visually inspected using the summary statistics method (ft_rejectvisual), trials contaminated by noise or artefacts (e.g., SQUID jumps, excessive noise) were interactively identified and rejected separately for EEG and MEG.

Subsequently, for both modalities an additional semi-automatic artefact detection was performed using z-value-based approach (ft_artifact_zvalue) to identify residual artefacts like muscle, or channel jumps in sensitive time ranges used for later event-related analyses (-0.2 to 0.6 s for localizer, 0.6 to 0.6 s for expectation task). Eye blinks were detected from the pupil size channel (band-pass filtered 1-15Hz) and respective trials removed if they occurred in pre-stimulus time range or up to 400ms after stimulus onset. The whole procedure led to on average 10 % rejected trials (range: 4 – 20 %). Artefact-cleaned trials were sorted into experimental conditions (localizer: orange, green, blue, magenta; colour expectations: additional split into colours expected and unexpected).

For visualization of the visually-evoked response (topographic maps, Fig. 3), condition-wise averaged event-related fields were computed per participant, low-pass filtered at 40 Hz using a zero-phase Butterworth filter, and subsequently entered into a grand average (ft_timelockgrandaverage) across participants.

**Figure 3.**
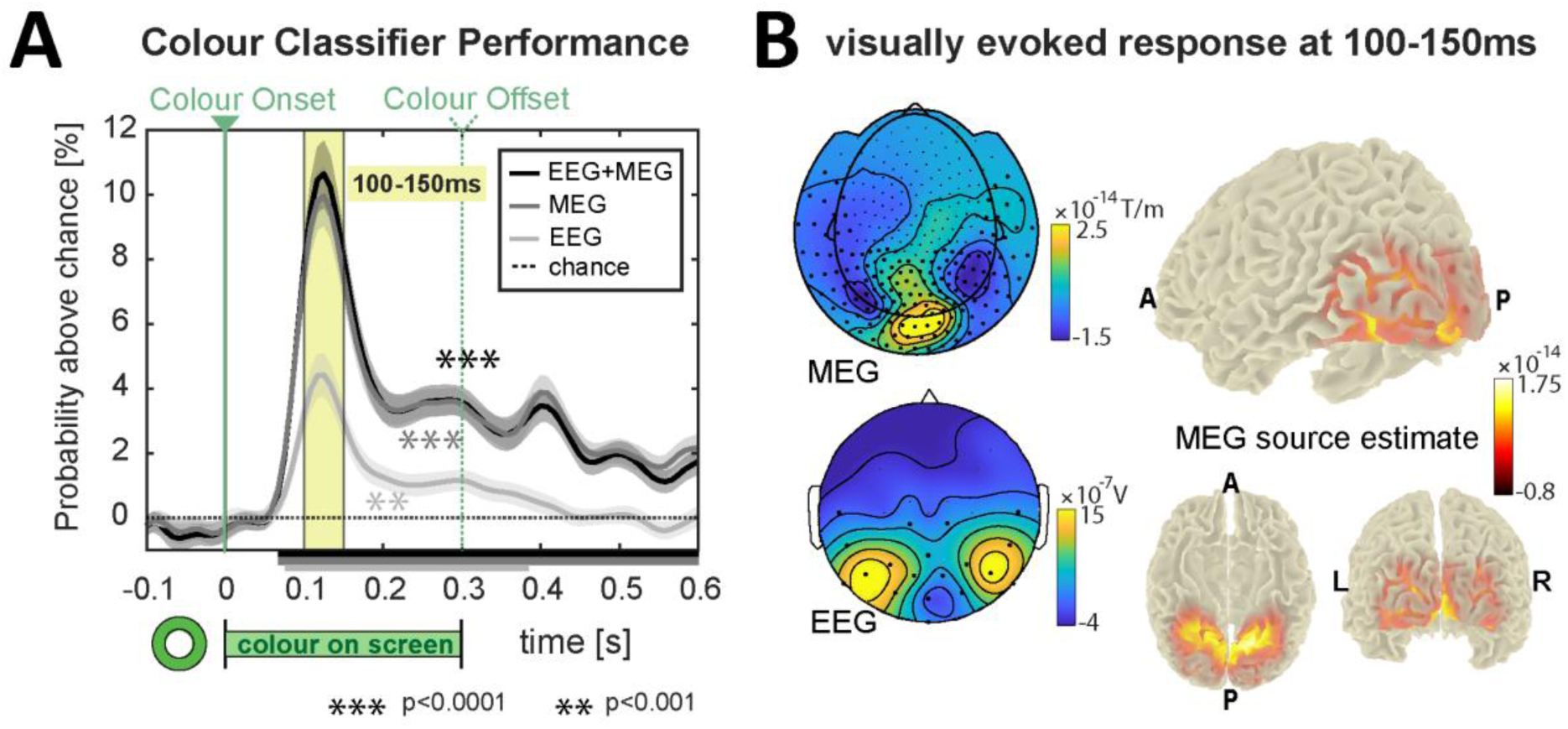
Reconstructing colour from localizer blocks. **A)** Probability of correct colour classification above chance level (25%) using EEG (light grey), MEG (dark grey), or the combination of both modalities (black) from posterior (occipito-temporo-parietal) sensors. All modalities show robust colour decoding beginning at ∼70 ms after stimulus onset (green vertical line), with MEG outperforming EEG and the combined modality yielding the highest accuracy at the decoding peak (100–150 ms). Shaded areas represent the standard error of the mean; horizontal bars indicate time windows of statistically significant decoding above chance level (25%, dotted horizontal line). Colour stimuli were presented between 0 and 300 ms, offset indicated by green dashed vertical line. **B)** Visually evoked responses during the time window of peak colour decoding (100-150ms). Event-related field maps on the left and an MEG-based source estimate on the right (LCMV Beamforming: left side, inferior and rear side view, letters demarcate A: anterior, P: posterior, L: left, R: right) reveal activation maxima in visual cortex, ranging from early visual areas to higher-level, ventral-extrastriate, colour-sensitive regions (presumably encompassing V1–V4 and VO1). Bold black dots in the field maps indicate sensor positions used for decoding.

#### MEG – source analysis

Source analysis was performed using FieldTrip toolbox in MATLAB (Oostenveld et al., 2011) (RRID:SCR_004849). Source estimates were based on thirty-six participants, one participant was excluded due to missing anatomical MRI data. Epoched data were baseline-corrected, low-pass filtered at 30 Hz, downsampled to 120 Hz (ft_resampledata), and restricted to 0–400 ms. Trials from all colour conditions were combined to compute time-locked responses and covariance matrices (ft_timelockanalysis). Structural MR images were realigned to the MEG coordinate system using fiducials (ft_volumerealign), and refined with digitized headshape data via an iterative closest point algorithm and visual inspection. MRIs were segmented into brain, skull, and scalp (ft_volumesegment), and a single-shell headmodel was created (ft_prepare_headmodel, method = ‘singleshell’). A subject-specific source grid (8 mm, warped from MNI template) was prepared (ft_prepare_sourcemodel). The MEG leadfield was computed using ft_prepare_leadfield with the individual headmodel, warped grid, and sensor geometry (reducerank = 2).

Source activity was estimated using an LCMV beamformer (ft_sourceanalysis) with unit-noise-gain normalization and fixed dipole orientation. Activity was reconstructed for 100–150 ms post-stimulus, quantified as the absolute dipole moment per voxel, and averaged across participants. Individual source maps were interpolated to a standard anatomical template and visualized (ft_sourceplot).

#### EEG / MEG – Colour decoding analyses

##### Sensor selection

Multivariate pattern analyses were restricted to posterior (occipital/ parietal/ temporal) sensors to target colour-related responses in visual cortex (Brouwer and Heeger, 2009; Hajonides et al., 2021; Rosenthal et al., 2021), yielding 154 sensors for MEG (all with “T”, “O”, “P”) and 19 electrodes for EEG (CP1, T7, P7, CP2, T8, P8, Pz, Oz, Iz, P3, PO7, PO9, PO3, O1, P4, PO8, PO10, PO4, O2), see markings in Fig. 3B.

##### Pre-processing

Data were low-pass filtered at 30Hz and downsampled to 120Hz (ft_resampledata). Data from all localizer and expectation task conditions were slightly cropped (localizer: -0.2-0.6 s, colour expectation: -0.6-0.6 s), and concatenated into a continuous dataset to perform normalization across the entire recording. Specifically, data were z-scored per channel across time and then segmented back into their original trial structure. Single-trial baseline correction was applied (localizer: −0.2 to −0.1 s before stimulus onset; expectation task: −0.2 to −0.1 s before outline onset). Data were converted into trial x channel x time matrices for subsequent decoding analyses.

##### Multivariate pattern analysis

Multivariate pattern analysis were conducted at the single participant level using custom MATLAB scripts and the MVPA-Light toolbox (Treder, 2020). To improve signal-to-noise ratio, time series were smoothed using a Gaussian kernel (*σ* = 25 ms) before decoding. Classification was performed using a multiclass linear discriminant analysis, yielding time-resolved estimates of class probabilities. Decoding results were then averaged across participants at the group level.

##### Colour Localizer

Analyses were performed separately for each modality (MEG, EEG, and combined MEG+EEG). Trial counts were balanced across colour conditions by random subsampling to match the minimum number of trials per condition. A k-fold cross-validation procedure (k = 5) was implemented and repeated 100 times using random partitioning of trials into folds (mv_classify_across_time). Decoding results were averaged across repetitions.

##### Colour Expectation Task

To retrieve sensory colour representations during the expectation task, classifiers were trained on colour localizer data and then tested on expectation task trials. Testing was performed both during the post-stimulus interval (stimulus decoding, Fig. 4) and the pre-stimulus interval (expectation template phase, no colour on screen, Fig. 5). For the main analyses (Fig. 4 and 5), expectation trials were pooled across conditions irrespective of validity to maximize signal-to-noise ratio. Trials were labelled according to either the perceived colour (stimulus decoding) or the expected colour (template decoding). To further enhance signal-to-noise ratio, subtrial averaging was performed by randomly grouping trials within each condition into sets of 10 and averaging within each group (Adam et al., 2020: “mini blocks”; Hebart et al., 2018: “supertrials”; Scrivener et al., 2023: “pseudotrials”). This procedure was repeated 100 times, each time followed by decoding, to account for variability introduced by random grouping. Final decoding performance was obtained by averaging across repetitions.

**Figure 4.**
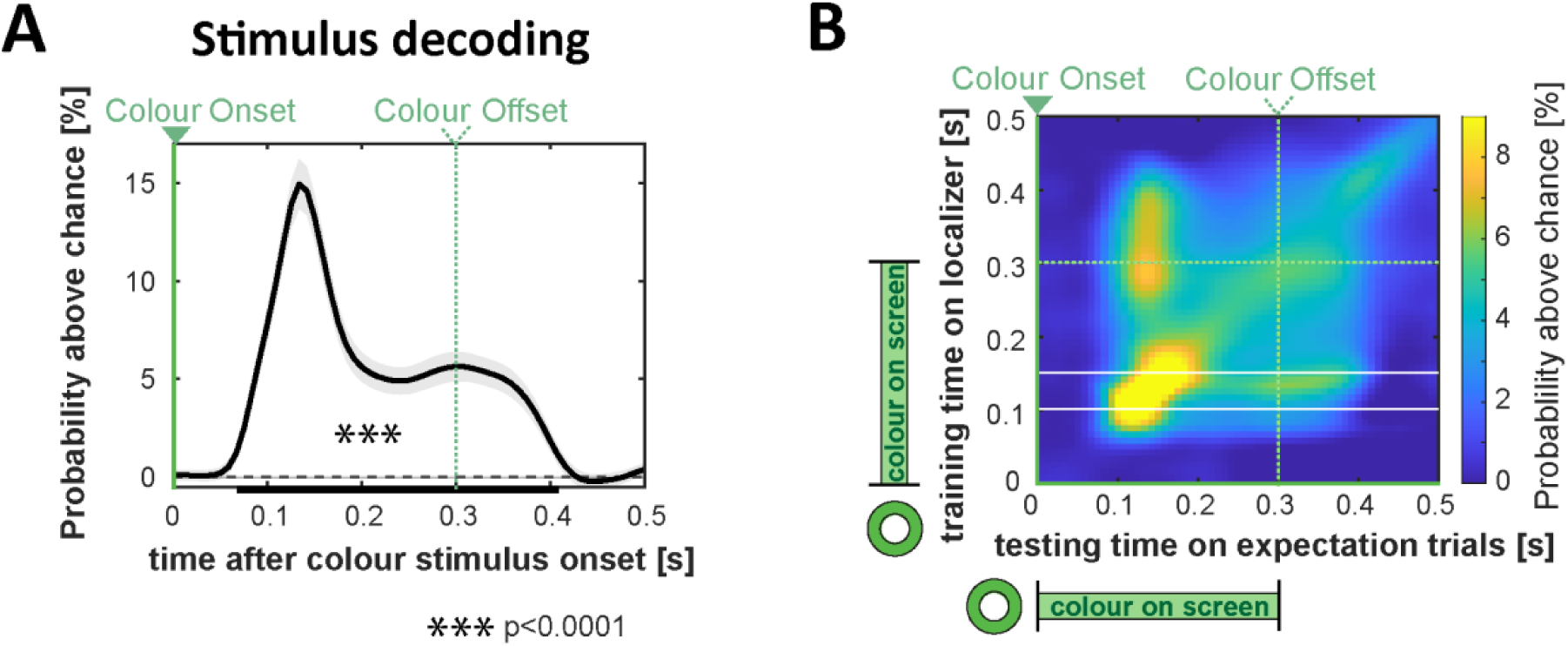
Decoding colour from expectation trials post-stimulus onset. **A)** Above-chance probability of colour classification when trained on the localizer signal averaged across the 100–150 ms window (white horizontal lines in B). The waveform shows significant above-chance decoding (chance = 25%, dashed line) from 67–408 ms after stimulus onset (p < 0.001, cluster-based permutation test), with a peak at 133 ms. Shaded areas denote the standard error of the mean (SEM), the black horizontal bar marks the significant time cluster. **B)** Cross-dataset temporal generalization matrix showing colour classification performance (probability of correct classification above chance) when training on individual time points from the localizer trials (y-axis) and testing across post-stimulus time points of the colour expectation trials (x-axis). Successful colour decoding is observed after stimulus onset (marked by solid green vertical line), peaking when both training and testing time points fall between 100–200 ms. This range closely aligns with the previously identified window of maximal localizer decoding performance (100–150 ms, indicated by white solid horizontal lines). Colour stimuli were presented between 0 and 300 ms for both localizer and expectation trials, offsets depicted by green dashed lines.

**Figure 5.**
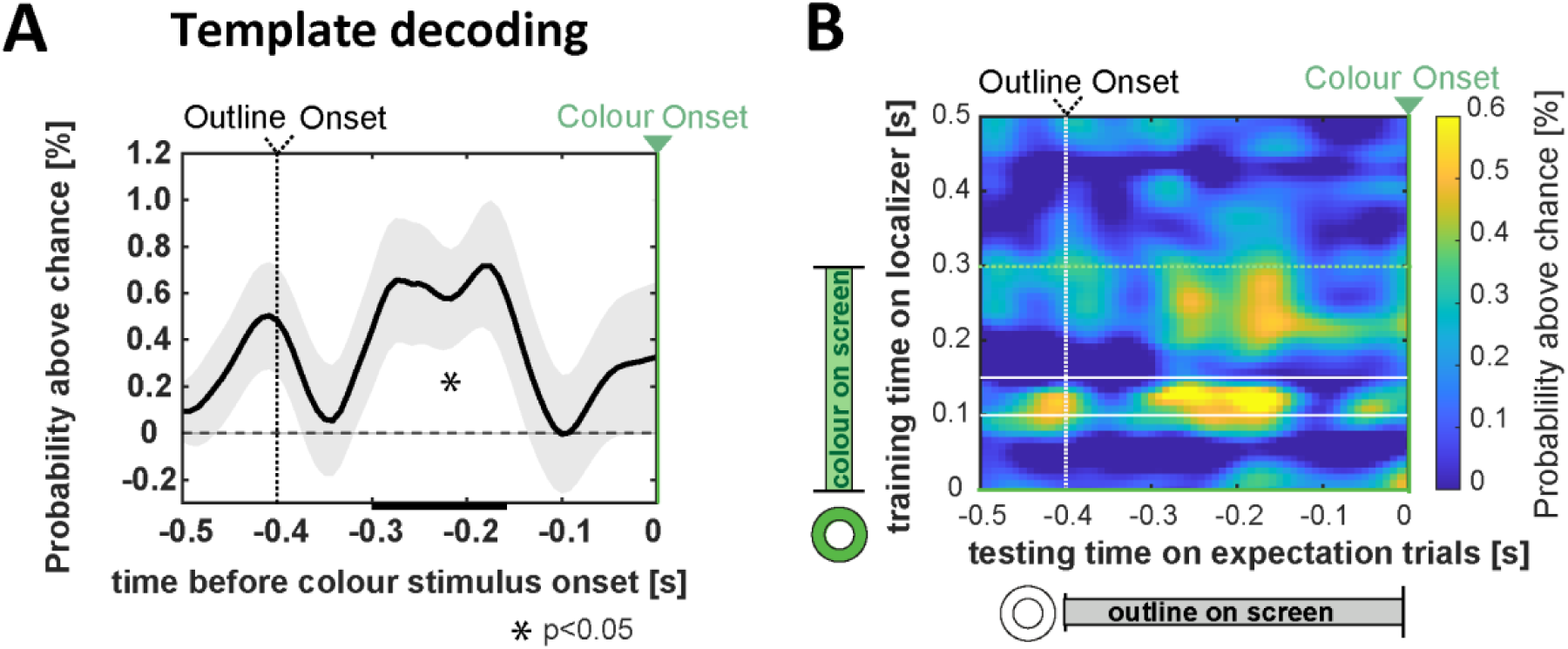
Decoding the colour template from expectation trials pre-stimulus onset. **A)** Above-chance probability of colour classification when trained on the localizer signal averaged across the 100–150 ms window (white solid lines in B). The resulting waveform shows significant above-chance decoding from -300 to -158 ms after the onset of the grey outline (p = 0.02, cluster-based permutation test). Shaded areas represent the standard error of the mean (SEM); the black horizontal bar marks the significant time cluster. Expectations emerging in the pre-stimulus period do, indeed, carry decodable sensory colour information. **B)** Cross-dataset temporal generalization matrix showing colour classification performance (probability of correct classification above chance) when training on individual time points from the localizer trials (y-axis) and testing across pre-stimulus time points of the colour expectation trials (x-axis). The grey outline was presented between -400 and 0 ms (-400ms corresponds to the onset of the outline, white vertical dotted line; 0 ms corresponds to the onset of the coloured stimulus, green vertical line). Horizontal green dashed line depicts colour offset in localizer trials. Peaks of highest colour decoding performance in the pre-stimulus time range align with the pre-defined training window of 100-150ms (white solid horizontal lines), reflecting early sensory colour representations.

##### Expected versus unexpected colours

To investigate potential influences of colour expectation on colour decoding after stimulus onset (see Supplementary Material, Figure S1), cross-dataset colour decoding (training on localizer and testing on expectation trials) was compared between trials with valid colour order (80% of trials) and those where an unexpected colour was presented (20% of trials). To equate trial numbers, random subsets of expected colours were drawn that matched the number of unexpected trials in the respective colour bin before decoding (subsets drawn 100 times, decoding results averaged). Given the reduced trial count (matched to the number of invalid trials), no subtrial averaging was performed for expectation trials.

##### Time × time generalization analysis

was conducted by training classifiers at each time point of the localizer data and testing them across all time points of the expectation task (*mv_classify_timextime*; Fig. 4 and 5 A, Fig. S1).

##### Time window of interest analysis

To optimize signal-to-noise ratio in the time window of interest (Fig. 4–5: localizer peak decoding: 100–150 ms, Fig 6: exploratory time window of enhanced responses to unexpected stimuli: 275-325ms), localizer data were averaged across this interval prior to training. The resulting model for this combined time range was then tested across all time points of the expectation task (Fig. 4 and 5 B, Fig. S1).

##### Statistical validation

Decoding results were statistically evaluated with custom scripts utilizing the nonparametric cluster-based permutation test *ft_timelockstatistics* implemented in the Fieldtrip toolbox (Oostenveld et al., 2011) (RRID:SCR_004849). Analyses were conducted across participants on time-resolved classifier outputs. Cluster-based permutation testing was performed using a dependent-samples t-statistic (observed vs. chance level). Temporal clusters were defined by grouping adjacent time points exceeding a cluster-forming threshold of p < 0.05. Cluster-level statistics were computed using the maximum sum of t-values within each cluster. Permutation distribution was obtained using a Monte Carlo procedure (10,000 randomizations). Statistical significance was assessed using a one-tailed test (testing for decoding performance above chance), with a cluster-level significance threshold of alpha = 0.05. To assess time ranges of statistical significance in cross-temporal generalization matrices, an analogue custom multi-dimensional version of the cluster-based permutation test (one-tailed, alpha = 0.05, cluster alpha = 0.05) was used to identify clusters in time x time space (Spaak, 2024).

To compare decoding performance across modalities for the localizer, a simple paired t-test was conducted for the average performance in the pre-defined time window (100-150ms), with alpha = 0.05.

## Results

### Behavioural performance

Participants demonstrated high accuracy in detecting luminance changes during the colour localizer blocks (hit rate = 95.9%, SEM = 0.85; false alarm rate = 0.76%, SEM = 0.18; reaction time = 448.3 ms, SEM = 6.12) and hue changes during the colour expectation blocks (hit rate = 96.03%, SEM = 0.78; false alarm rate = 0.53%, SEM = 0.13; reaction time = 625.8 ms, SEM = 6.98). Performance in the colour expectation blocks was near ceiling for both expected and unexpected colour sequence trials, with no significant differences in hit rate (expected: 96.18%, unexpected: 95.47%, t(36) = 1.18, p = 0.24, d = 0.19) or false alarm rate (expected: 0.53%, unexpected: 0.54%, t(36) = -0.21, p = 0.84, d = -0.034). However, colour expectation exerted a small but significant influence on reaction times: responses were slightly faster when colours appeared in the expected rather than the unexpected sequence (expected: 624.7 ms, SEM = 6.86; unexpected: 630.0 ms, SEM = 7.74; difference = 5.33 ms, t(36) = -2.22, p = 0.033, d = -0.36; see Figure 2). Note that because colour sequence was task-irrelevant, only modest behavioural effects were expected. Nevertheless, the behavioural results indicate that hue changes were, indeed, detected more rapidly when they occurred on an annulus of the anticipated colour, speaking for the behavioural relevance of colour expectations elicited by colour order.

### Decoding colour from the brain response – Colour localizer task

Colour could be reliably classified from neural activity from posterior EEG and MEG sensors during the localizer task, beginning 67 ms after the onset of the coloured annulus (Figure 3A; p < 0.0001, cluster-based permutation test). Classification accuracy peaked around 125 ms for both modalities, with MEG outperforming EEG (note that EEG included only 19 sensors, whereas MEG used 154). The highest decoding performance above chance was achieved when EEG and MEG data were combined (EEG: +4.4%; MEG: +10.0%; combined: +10.6%), suggesting that the individual modalities both contributed unique variance. A time-resolved comparison of classification accuracy revealed a significant advantage of MEG over EEG across the entire time window from 75 ms onward (p < 0.0001, cluster-based permutation test). However, within the peak interval of classification performance (100–150 ms), the combination of EEG and MEG significantly outperformed MEG alone (paired t-test, p = 0.0304). Based on this, we used the combined EEG–MEG signal for all subsequent analyses.

Examining visually evoked responses during the peak decoding window (100–150 ms), we observed activation maxima in early visual cortex extending into ventral occipital regions, likely including areas V1–V4 up to VO1 (Figure 3B). This pattern is consistent with prior reports that these regions encode colour information (Brouwer and Heeger, 2013, 2009). Notably, the time window also overlaps with the P1 and N1 components of the visual evoked response, which have been associated with spatial and feature-specific encoding -including colour- during attentional selection (Anllo-Vento et al., 1998; Mangun and Hillyard, 1991; Moher et al., 2014; Zhang and Luck, 2009).

Interestingly, colour information could still be decoded after stimulus offset at 300 ms. This persistence may reflect ongoing cortical representations of perceptual colour information, but it could also indicate the decoding of transformed signals (e.g., offset responses or memory-related activity). To ensure that decoding primarily reflected sensory rather than post-perceptual processes, subsequent analyses specifically focused on the early time window (100–150 ms)—the period of peak classification accuracy—for training the decoding model.

In the next step, we used the colour decoding model trained on the localizer blocks to quantify colour information in the activity patterns during the expectation task trials.

### Early sensory representation of colour after stimulus onset – Expectation task

We trained a decoding model on the localizer signal averaged across its peak decoding window (100–150 ms) and tested it on the post-stimulus onset period in the expectation trials (0–500 ms; see Figure 4A). This cross-decoding analysis revealed significant colour decoding in the expectation trials between 67–408 ms following the onset of the coloured stimulus (p = 0.0001, cluster-based permutation test), with maximal decoding at 133 ms (+14.9% above chance level). This demonstrates that (1) sensory colour information is represented similarly between localizer and expectation tasks, as the expectation trials could be cross-decoded from the localizer data, and (2) this information is strongest during an early stage of the visually evoked response, corresponding to early sensory feature processing. As expected, sensory colour information was present in visual cortex shortly after stimulus onset. Notably, this information persisted for ∼100 ms beyond the physical presence of the stimulus on the screen (see Figure 4A, dashed vertical green line). It is likely that colour representations in early visual areas were subsequently overwritten by visually evoked responses to the stimulus offset, which did not align with the original sensory colour information.

In a follow-up analysis, we trained decoding models on the colour localizer data separately at each time point within the 0–500 ms window and tested them across all post-stimulus onset time points in the colour expectation trials (0–500 ms; see Figure 4B) – using a cross-temporal generalization method (King and Dehaene, 2014; Meyers et al., 2008). A prominent peak in classification accuracy emerged when both training and testing time points fell between 100–200 ms, which confirms that cross-decoded colour information is primarily driven by early sensory processing and dovetails with our choice of the 100–150 ms localizer peak as the main training interval (white horizontal lines in the matrix). The cross-temporal generalization matrix additionally revealed residual colour decoding beyond 400ms, but only when training time points corresponded to the offset period of the localizer (diagonal of the matrix). This is consistent with the interpretation that after stimulus offset, colour information does not completely disappear but changes into a representational format that no longer corresponds to the early sensory-driven response.

Finally, an additional analysis comparing post-stimulus onset colour decoding between valid and invalid colour order trials for the 100-150ms training window revealed no significant differences (cluster-based permutation test, 0–500 ms: –1.68 < all t < 0.78), suggesting that the amount of decodable sensory colour information after stimulus onset was unaffected by whether the colour was expected or not. We also performed complete time–generalization analyses separately for expected and unexpected colours. These point towards an increase in decodability for unexpected colours in a later time-window around ∼300 ms post-stimulus (training and testing), which would be consistent with previous reports that surprising stimuli elicit stronger visual cortical responses (Kok et al., 2016, 2012; Meyer and Olson, 2011; Richter et al., 2018). The respective analyses can be found in the Supplementary Material (Figure S1).

### Predictive early sensory colour template before stimulus onset – Expectation task

After establishing robust colour decoding in the post-stimulus period, we next examined whether sensory colour information was already present during the pre-stimulus phase—when no colour was physically displayed—potentially reflecting internally generated expectation templates.

Corroborating our hypothesis, we found evidence for a predictive colour template in the pre-stimulus time window (Figure 5A). When training on the averaged localizer signal between 100– 150 ms and testing across the pre-stimulus interval starting at the onset of the grey placeholder outline (–400 to 0 ms), significant above-chance classification emerged between –300 and –158 ms in the template time range before stimulus onset (black horizontal bar; p = 0.02, cluster-based permutation test, Figure 5A). This pre-stimulus neural representation of expected colour information cannot be explained by physical input or trial-by-trial priming. Instead, we argue that it reflects an internal colour expectation template shaped by the statistical regularities of the colour sequence. Importantly, significant decoding was visible ∼100 ms after the onset of the grey placeholder, consistent with a “ping approach,” in which the visually evoked response to a neutral stimulus -here the grey placeholder- is modulated by the expected colour information that can uncover otherwise activity-silent internal representations (Kandemir et al., 2024; Wolff et al., 2017, 2015).

For completeness we performed again a follow-up cross-temporal generalization analysis (Fig. 5B, training on each time point of the localizer and testing on all time points of the pre-stimulus interval). As evident in the resulting cross-decoding matrix, activity in the previously chosen peak localizer window (100-150 ms, white horizontal lines) is, indeed, predominantly driving the decoding of the predictive template in the expectation task. In other words, cross-decoded colour information in the expectation time range primarily reflects early sensory colour representations.

## Discussion

Here, we investigated whether expectations give rise to predictive colourful templates in early visual cortical areas. We found that predictable colour sequences induced reliable expectations, with decodable colour information emerging in early visual cortex before stimulus onset. After stimulus onset, decoding of sensory colour information was not modulated by expectation. However, unexpected colours slowed behavioural responses and seemed to be associated with enhanced late neural activity – a pattern consistent with the generation of prediction error signals.

### Decoding colour from the brain response to coloured stimuli

Consistent with previous research, colour information could be decoded from cortical responses to coloured stimuli in early visual areas. Significant decoding emerged as early as 70 ms after stimulus onset, peaked around 120 ms, and gradually declined, with colour information persisting beyond stimulus offset (Bocincova and Johnson, 2019; Hajonides et al., 2021; Hermann et al., 2022; Sandhaeger et al., 2019; Sutterer et al., 2021). Similar persistence of colour decoding beyond stimulus offset has been reported during the encoding phase of visual working memory (Bae and Chen, 2024; Bocincova and Johnson, 2019; Che et al., 2022; Hajonides et al., 2021). Although in the present paradigm, participants had no incentive to actively maintain colour information in memory, perceptual colour information persisted for approximately 100 ms after stimulus offset. Afterwards, the representation appeared to transform keeping colour identity but no longer matching sensory colour information (Fig. 4B). Since our goal was to identify sensory representations rather than post-perceptual processes, we trained the colour classifier on the initial decoding peak (100–150 ms after stimulus onset) to search for predictive colour templates.

### Expectations contain sensory colour information

Because expectations are inherently memory-based, an important question is the extent to which expectation templates resemble perceptual colour representations. For orientation stimuli, representations maintained in working memory have been shown to take a more abstract form (Duan and Curtis, 2024; Kwak and Curtis, 2022), or gradually shift toward more categorical formats along the visual hierarchy (Chunharas et al., 2025). For colour stimuli, representations maintained in working memory have been reported to resemble both perceptual and categorical codes with a preference for categorical representations arising during the memory delay already at the level of V4 and VO1 (Yan et al., 2023). Although the colours used in the present study (orange, blue, magenta, green), were distinct both perceptually and categorically (Berlin and Kay, 1969; Brouwer and Heeger, 2013; Rosenthal et al., 2021), pre-stimulus cross-decoding peaked when classifiers were trained on early sensory responses (100–150 ms; Fig. 5B). This suggests that expectation templates more closely resembled early sensory colour representations than more abstract or categorical codes, which would be expected to emerge at later stages of colour discrimination, or after stimulus offset. Consistent with this interpretation, Bartsch et al. (2017) found that colour-selective responses at 110-150ms primarily reflect task-independent sensory colour differences, whereas category-like sharpening emerged later at 275-390ms, and only during active colour discrimination, which was discouraged in the present localizer task. Future studies comparing within- and between-category colour expectations could further clarify the representational format and resolution of predictive colour templates.

### Eliciting colour expectations through sequential structure

A key feature of this study was the use of colour order to elicit expectations. The expected colour was neither presented earlier within the same trial, as typical in working memory designs (Bae and Chen, 2024; Bocincova and Johnson, 2019; Hajonides et al., 2021), nor cued via another sensory modality (e.g., auditory cues), as in prior work on expectation templates (Kok et al., 2017, 2014). Because colour order was task-irrelevant, expectation decoding was dissociated from effects of attention and response preparation. Combined with an independent perceptual colour localizer and cross-decoding, this design ensured that successful decoding reflected internal colour representations elicited by learned statistical regularities, rather than priming, sensory adaptation, or task-related strategies such as colour naming. Under these conditions, the emergence of a decodable representation of the upcoming colour is particularly notable, especially because the preceding bottom-up input was a different colour.

A central question in the literature is whether expectation effects require conscious attention or instead reflect more automatic processes. Although participants were informed about the colour order, most reported not actively attending to it, or doing so only briefly before disengaging; some even concluded after encountering invalid trials that no consistent sequence existed. Nevertheless, expected colours elicited faster responses, indicating that explicit tracking of the sequence was not required for expectations to influence behaviour, consistent with more implicit predictive mechanisms. This is in line with prior findings showing that prediction effects depend on attending the stimuli (Richter and De Lange, 2019) but not the predictive structure (Richter et al., 2018), or the predicted feature itself (Kok et al., 2017). Still, expectation templates may have been stronger had participants explicitly attended the colour order.

Recent work has suggested that feature-specific preactivations in predictable sequence paradigms may reflect classifier confounds rather than genuine predictions (Abdoun et al., 2026, but see Demarchi et al., 2026). Specifically, prediction-template effects in tone sequences may arise because neighbouring tones are more similar in frequency and therefore more likely to be confused by the classifier. This explanation is unlikely here because sequence order was randomized across participants and reversed halfway through the experiment, eliminating any potential influence of systematic similarity between neighbouring colours.

### Decoding colour from grey stimuli

The only visual input during the expectation interval was a uniform grey placeholder. Unlike prior work that decoded implied colour from greyscale objects, e.g., the yellow of a grey banana, which rely on object–colour associations stored in long-term memory (Bannert and Bartels, 2013; Retter et al., 2023; Teichmann et al., 2019; Zhao et al., 2024), the placeholder was identical across trials and lacked any intrinsic colour association. Accordingly, successful decoding must reflect the already expected upcoming colour and not colour implied by the placeholder itself. Consistent with this interpretation, decoding emerged approximately 100 ms after placeholder onset, which matches the timing of responses to physically presented colours rather than the delayed dynamics typically observed for implied colour (Teichmann et al., 2019).

Although the placeholder was originally introduced to provide a temporal anchor that supports the allocation of attention to the upcoming stimulus (Bartsch et al., 2018; Jefferies and Di Lollo, 2015; Wildegger et al., 2017; Woodman et al., 2009), it may also have functioned analogously to an “impulse” or “ping” used to probe latent neural representations (Kandemir et al., 2024; Wolff et al., 2017, 2015). Hence, the grey placeholder may have actually served a dual function: beyond marking the temporal task structure, the elicited visual response might have supported decoding of pre-activated expected colour information.

### Responses to expected versus unexpected colours

Unexpected colours (20% trials) elicited slightly slower responses. However, expectation did not modulate early sensory colour representations: classifiers trained on the 100–150 ms localizer response decoded expected and unexpected colours equally well (cluster-based permutation test, 0–500 ms after stimulus onset: –1.68 < all t < 0.78). This finding dovetails with previous studies showing that colour congruency effects do not influence early stages of visual processing but emerge typically after 200 ms (Proverbio et al., 2004; Teichmann et al., 2020). Exploratory analyses (Suppl. Figure S1) revealed, indeed, stronger neural responses to unexpected colours in the P300 time range that were associated with slower responses - a pattern consistent with prediction error signals and previous predictive processing studies (Kok et al., 2016, 2012; Meyer and Olson, 2011; Richter et al., 2018). Still, the exploratory nature of these findings warrants caution, and future research will be needed to anchor these findings with more certainty.

### Relation to attention-related preparatory activity

Although colour predictions were intentionally task-irrelevant, our findings relate to research on attentional preparation to specific features or objects. Attentional preparatory activity can spread globally during the pre-stimulus period (Menceloglu et al., 2026), and recruit feature-specific (e.g., colour- or motion-selective) visual areas (Chawla et al., 1999; Shibata et al., 2008). However, whether this activity reflects sensory representations or abstract templates remains debated. While some studies report that preparatory attention templates contain object-specific representations (like letters, people, or cars) during visual search (Gayet and Peelen, 2022; Peelen and Kastner, 2011; Stokes et al., 2009), a recent fMRI study reported that basic feature representations do not generalize to pre-stimulus preparatory activity, concluding that preparatory attention relies primarily on non-sensory representations (Gong et al., 2022). The latter differs from our results, possibly because their localizer (single motion direction) differed from task stimuli (overlaid motion directions), potentially limiting cross-task generalization. Moreover, no impulse stimulus was used, which may be important for revealing otherwise activity-silent sensory responses (Kandemir et al., 2024).

### Conclusion

We find that predictable colour sequences lead the brain to form sensory colour templates in early visual cortex before stimulus onset. Our findings contribute to the ongoing debate regarding the nature of preparatory signals and suggest that pre-stimulus expectations carry sensory colour information. This extends previous work on anticipatory orientation templates (Kok et al., 2017), and demonstrates that predictive mechanisms can pre-activate diverse basic visual features across spatial and non-spatial domains, including colour.

## Acknowledgements

We thank Vivian Vriezekolk for support with data collection. This project has received funding from the European Union’s Horizon Europe research and innovation programme under the Marie Skłodowska-Curie grant agreement No. 101061920 (Project Name: COLOUR) (MVB).

## Data and code availability

All data and code used for the analyses will be openly available (upon user registration) on the Radboud Data Repository Site (https://data.ru.nl).

## Author contribution

MVB: Conceptualization, Methodology, Software, Analysis, Investigation, Writing – Original Draft preparation, Visualization, Project Administration, Funding Acquisition. ES: Methodology, Software, Writing – Review and Editing. FDL: Conceptualization, Methodology, Resources, Writing – Review and Editing, Supervision, Project Administration, Funding Acquisition

## Supplementary information

Supplementary Material, contains Figure S1.

