## Supplementary Material for "Colourful predictive templates in early visual cortex"

### Supplementary analysis: Decoding colour from expected versus unexpected trials – post-stimulus onset

We performed time–generalization analyses of expected versus unexpected colours across all post-stimulus training and testing time points to investigate potential influences of color expectation on colour decoding after stimulus onset. Because colour order was violated in only 20% of trials, decoding for unexpected colours was compared with equally sized subsets of expected trials (subsets drawn 100 times; decoding results averaged). Across the full cross-temporal generalization matrices, there were no significant differences in stimulus colour decodability between expected and unexpected trials (cluster-based permutation test across full cross-temporal generalization matrices $p > 0.05$).

However, post-hoc visual inspection (Suppl. Figure S1) suggested a potential increase in decodability for unexpected trials around ~300 ms post-stimulus (training and testing), consistent with previous reports that surprising stimuli elicit stronger visual cortical responses (Kok et al., 2016, 2012; Meyer and Olson, 2011; Richter et al., 2018). An exploratory analysis of this time window further supported this observation: training on a 50 ms window around 300 ms (averaged signal) revealed higher decoding performance for unexpected trials between 300–367 ms after stimulus onset (mean probability above chance: $3.28 \pm 0.81$, 95% CI, unexpected; $2.44 \pm 0.57$, 95% CI, expected; Suppl. Figure S1C). Given that we observed slightly slower response on unexpected colour trials (Figure 2B), this result is consistent with the notion that stronger neural representations of the unexpected colour in the P300 time range may be associated with slower reaction times on those trials. The effect also appeared to interact with behaviour, such that especially participants that do show the behavioural benefit for expected compared to unexpected colours exhibited greater decoding scores for unexpected trials (Suppl. Figure 1D).

Although these findings align with the idea that a strong neural representation of an unexpected colour (prediction error) interferes with rapid reaction to hue changes, they are post-hoc and exploratory. Therefore, we do not interpret them further, but rather suggest them as interesting avenues for future work.

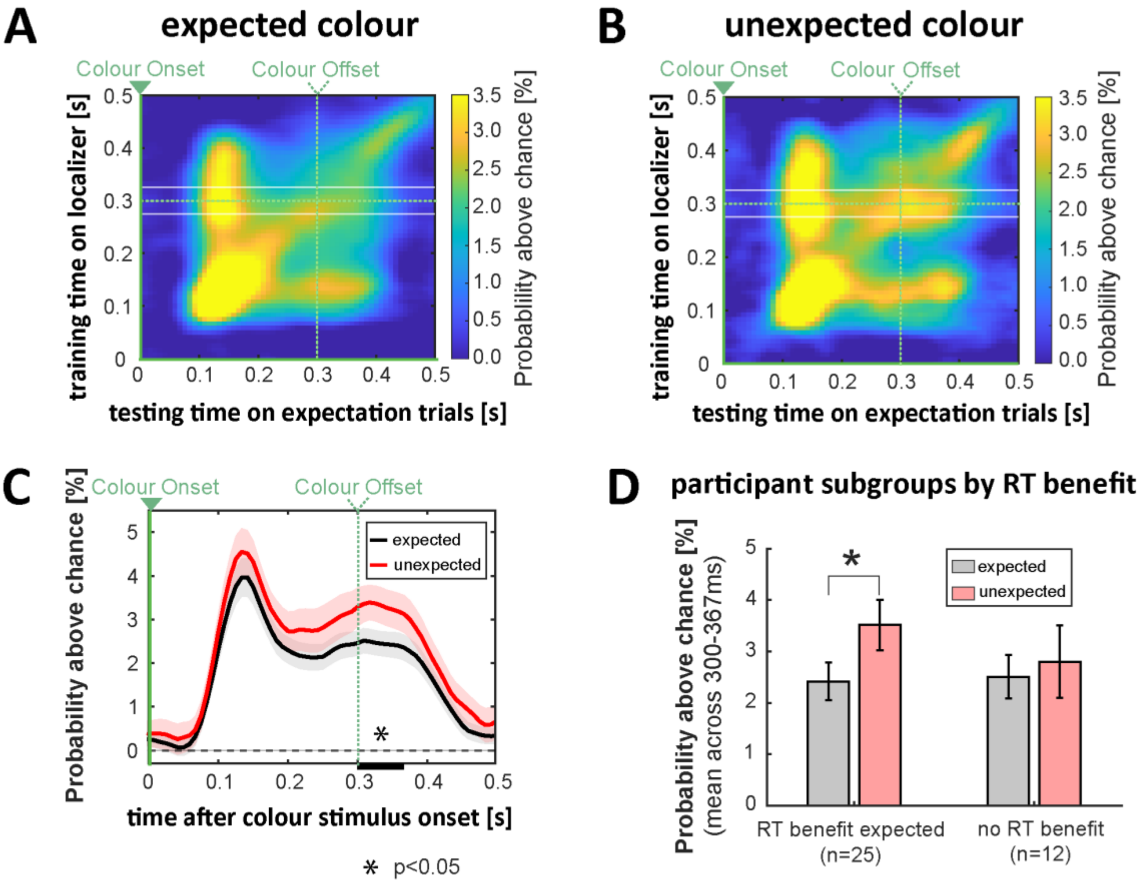


***Suppl. Figure S1. Decoding the colour from expectation trials post-stimulus onset: colour expected versus unexpected.*** *Cross-dataset temporal generalization matrix showing colour classification performance (probability of correct classification above chance) when training on individual time points from the localizer trials (y-axis) and testing across pre-stimulus time points of the colour expectation trials (x-axis) for trials where* ***A)*** *the expected colour was presented,* ***B)*** *an unexpected colour was presented (violation of colour order on 20% of trials). A visual inspection indicates increased decoding performance for unexpected colours, especially in the later P300 time-range (training and testing time).* ***C)*** *An exploratory analysis of the 50ms window around 300ms with the localizer trained on the averaged signal between 275–325ms, white solid lines in A) and B), shows higher probability of colour classification for unexpected colours (significant time cluster from 300 to 367 ms, marked by black horizontal bar). Shaded areas represent the standard error of the mean (SEM).* ***D)*** *Grouping participants based on whether they showed a reaction-time benefit for expected colours when detecting hue changes. Stronger neural representations of the unexpected colour in the P300 time range were observed particularly in participants who exhibited slowed reaction times on those trials*
